# Paradoxical Activation of GCN2 by ATP-competitive inhibitors via allosteric activation and autophosphorylation

**DOI:** 10.1101/2024.08.14.606984

**Authors:** Graham Neill, Vanesa Vinciauskaite, Marilyn Paul, Rebecca Gilley, Simon J. Cook, Glenn R. Masson

**Affiliations:** Division of Cellular and Systems Medicine, School of Medicine, University of Dundee, Dundee, UK; Drug Discovery Unit, Wellcome Centre for Anti-Infectives Research, Division of Biological Chemistry, University of Dundee, Dundee DD1 5EH, United Kingdom; Signalling Programme, The Babraham Institute, Babraham Research Campus, Cambridge CB22 3AT, U.K

## Abstract

Recently it has been found that General Control Non-derepressible 2 (GCN2) can be activated by an array of small molecule ATP-competitive inhibitors, including clinically relevant compounds such as Ponatinib, and compounds specifically designed to be GCN2 inhibitors, such as GCN2iB. Furthermore, we recently showed that GCN2 can be activated in cells by clinically approved small molecule RAF inhibitors. GCN2 is a drug target, specifically in cancers such as mesothelioma, and a better understanding of this paradoxical activation is required to develop drugs which truly inhibit the enzyme. Using biochemical assays and structural mass spectrometry, we present a model for how GCN2 is activated by these compounds by promoting an active conformation in the HisRS domain while competitively inhibiting the kinase domain. This conformation promotes activating phosphorylation of GCN2, potentially through phosphorylation of other activated GCN2 molecules which are not bound to compound. Together this model suggests that efforts to inhibit GCN2 would benefit from exploring allosteric routes rather than targeting the ATP-binding pocket of the kinase domain.

## Introduction

The Integrated Stress Response (ISR) is a crucial stress sensing pathway which protects cells from both external and internal sources of stress. Central to this pathway is the phosphorylation of the α subunit of the eukaryotic initiation factor 2 (eIF2), which results in a global depression in protein synthesis while facilitating the selective transcription of metabolism reprogramming genes which attempt to restore homeostasis ^1,2^. In higher eukaryotes, there are four obligate homodimeric eIF2α kinases (GCN2, PKR, PERK, and HRI), all of which sense a unique subset of stresses, become activated, autophosphorylate and subsequently phosphorylate eIF2α on serine 51, triggering the ISR. Given the pivotal role that these kinases have in sensing stress, it is unsurprising that their activity has been found to play a role in several diseases, not least cancer ^3–8^, which in turn has led to efforts to generate small molecule inhibitors targeting these kinases ^9–12^.

Activation of GCN2 is complex, requires domain reorganisation, and can occur through multiple diverse routes ^13,14^. The current prevailing model is that GCN2 can be activated by amino acid deprivation through either the increased rate of ribosomal collisions or the increased concentration of deacylated tRNA. The ribosomal collision route is thought to be dependent on formation of a complex of the ribosomal collision sensing protein GCN1 ^15–17^, the ribosomal P-Stalk ^18–21^, and the kinase Zak-alpha ^22^, while the tRNA-binding route is simpler, with GCN2 binding directly to deacylated tRNA^13^ – recent elegant work by Misra *et al.* has shown that both these routes of activation co-exist and allow GCN2 to respond to different stress sources.

Several recent studies have shown that within a cellular context ATP-competitive inhibitors, even those designed specially to inhibit GCN2, can paradoxically activate GCN2 and the ISR ^23–25^. Furthermore, several clinically relevant drugs have also been shown to activate GCN2 *in vivo* ^26,27^, raising clinically relevant questions on drug mechanisms. Typically, these compounds are inhibitory at high concentrations (> 1 µM), but at lower, approximately nanomolar concentrations, they facilitate GCN2 activation. Previously, a mechanism of how these drugs may facilitate activation was proposed whereby occupancy of compound in the kinase domain in only one of the monomers of GCN2 caused an allosteric activation of the neighbouring monomer ^25^.

We recently demonstrated that eight BRAF inhibitors, including all clinically approved drugs used in the treatment of melanoma (Dabrafenib, Encorafenib, Vemurafenib) also exhibit paradoxical activation/inhibition of GCN2. In this study we use purified recombinant human GCN2 and found that there was a disconnect between substrate (i.e., eIF2α) phosphorylation and auto-phosphorylation of GCN2, something that is not predicted in current models of GCN2 activation. To further investigate this, we used Hydrogen Deuterium Exchange Mass Spectrometry (HDX-MS) to determine that these compounds, alongside GCN2iB, bind as expected within the active site of the kinase domain of GCN2, but also induce a series of allosteric rearrangements of GCN2, which may account for its activation. Finally, by using a kinase dead mutant of GCN2 (K619R) we show that dimers of GCN2 are capable of phosphorylating other dimers of GCN2 in *trans*, and this is promoted by compound binding. Overall, this presents a new model of how these compounds may facilitate activation of GCN2, which accounts for the low concentration dependency and disconnect between autophosphorylation and substrate phosphorylation.

## Results

### Activation of GCN2 by Kinase Inhibitors produces a characteristic disconnect between kinase autophosphorylation and eIF2α phosphorylation

Recently, it has been shown that GCN2 may be activated via ATP-competitive small molecules such as GCN2iB – however, these data derive from cellular experiments, or *in vitro* experiments using fragments of GCN2 ^23–25^. For these studies, we used three RAFi that activate GCN2 and the ISR in cells (Dabrafenib, Encorafenib and LY-3009120) and one that fails (GDC-0879) and made comparisons to GCN2iB in a recombinant purified assay using both full length GCN2 enzyme and full length eIF2α substrate. Dabrafenib, Encorafenib, LY-3009120 and GCN2iB activated GCN2 (observed as an increase in the autophosphorylation at p-T899) at low concentrations whereas at high concentrations we observed inhibition (Fig 1A & 1B). In contrast, GDC-0879 had no effect on GCN2 activation. Furthermore, there was an apparent disconnect between the increased levels of GCN2 autophosphorylation (as measured by phosphorylation of T899) and subsequent substrate phosphorylation on eIF2α (as measured by phosphorylation of S51). This disconnect between p-GCN2 and p-eIF2α was not observed when GCN2 was activated by a higher (5x) ATP concentration which caused an increase in both p-eIF2α and p-GCN2 levels.

**Figure 1:**
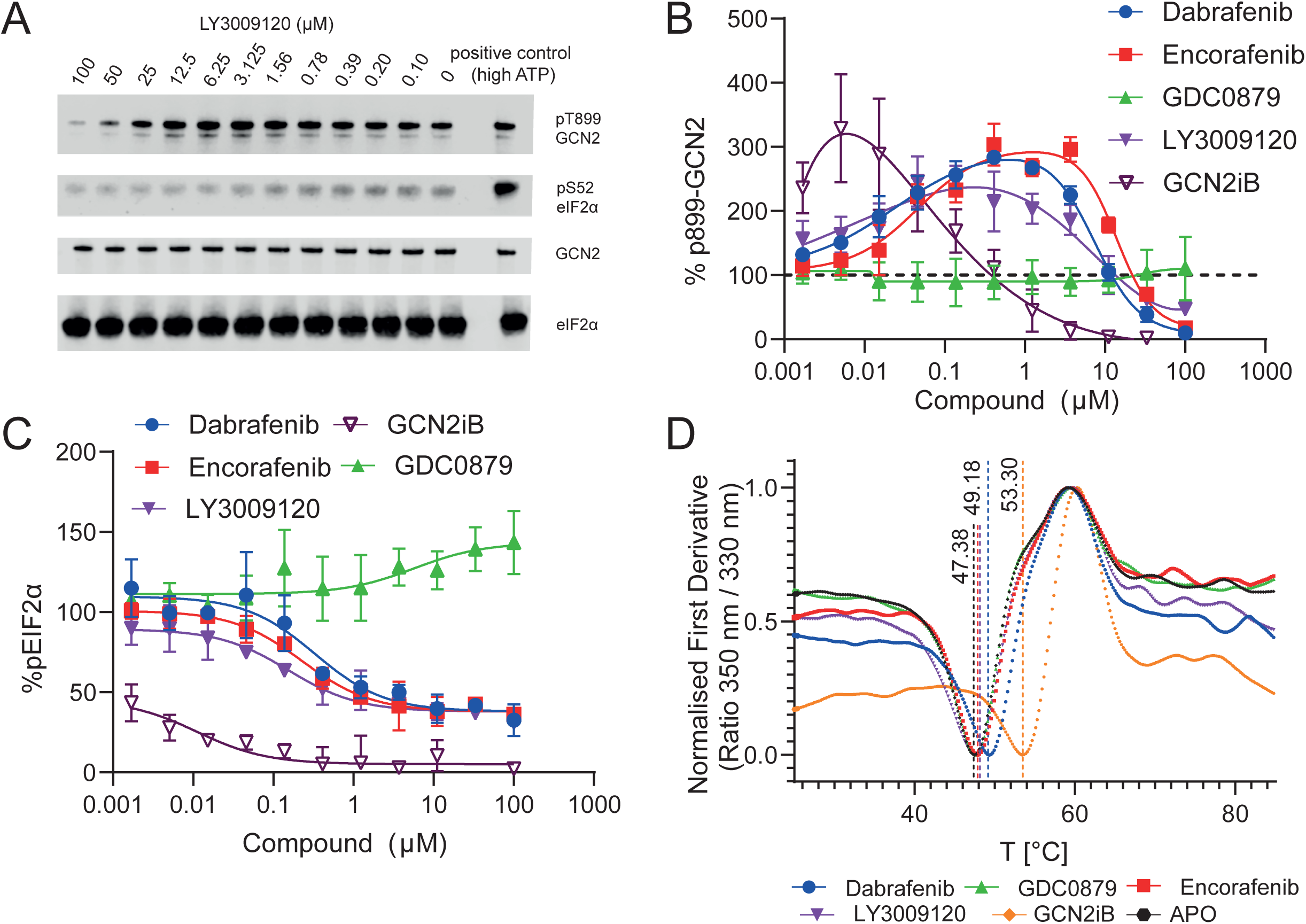
RAFi and GCN2iB promote GCN2 autophosphorylation, but not eIF2α phosphorylation. **(A)** Activity assay of human recombinant GCN2 phosphorylating eIF2α against a concentration gradient of LY3009120. **(B)** Quantified assays against 5 compounds – LY3009120, Dabrafenib, Encorafenib, GDC0879 and GCN2iB. pThr899 GCN2 levels peak at ∼1 µM for Dabrafenib, Encorafenib and GDC0879, and 10 nM for GCN2iB. GDC0879 does not appear to alter GCN2 activity **(C)** pS51 EIF2α levels across a gradient of compounds. **(D)** Thermal Unfolding assays of GCN2 against compounds. Lines represent nadirs of the first inflection point.

### Activation of GCN2 occurs at concentrations significantly below binding constants

The bell-shaped activation curves for GCN2 by RAFi and GCN2iB may suggest multiple binding sites, or different modes of binding. To investigate compound binding, we made use of Thermal Unfolding analysis and Surface Plasmon Resonance (SPR) (see Figure 1A-D, Supplementary Figure 1). Using SPR we found that Dabrafenib, Encorafenib, and LY3009120 all had micromolar binding affinities, which is consistent with the IC_50_ values observed on the “down-slope” of the activity assays. Further binding analysis using thermal unfolding, analysis of GCN2 demonstrated a wide range of compound stabilisation effects ranging from +0.6 °C (LY3009120) to +5.9 °C for GCN2iB. As per our activity assays, we did not observe binding for GDC-0879 in either binding assay.

### Dabrafenib, Encorafenib, and GCN2iB binding causes allosteric activation of GCN2

To assess the potential structural changes that accompany compound binding, we conducted Hydrogen Deuterium Exchange Mass Spectrometry (HDX-MS) on full-length human GCN2 in the presence of GCN2iB (see table 1), Dabrafenib and Encorafenib, and as a negative control, GDC-0879. To provide a better three-dimensional context for these HDX-MS data, we created an AlphaFold/ColabFold ^28,29^ model of human dimeric GCN2 which fulfilled many of the known intramolecular interactions which have been observed previously ^30–32^, including the recently published structure of the human HisRS-like domain of GCN2^33^ (Figure 2B). Mapping HDX-MS data onto this structure we found a good degree of agreement against likely folded sections, loops and areas of disorder (Figure 2C).

**Figure 2:**
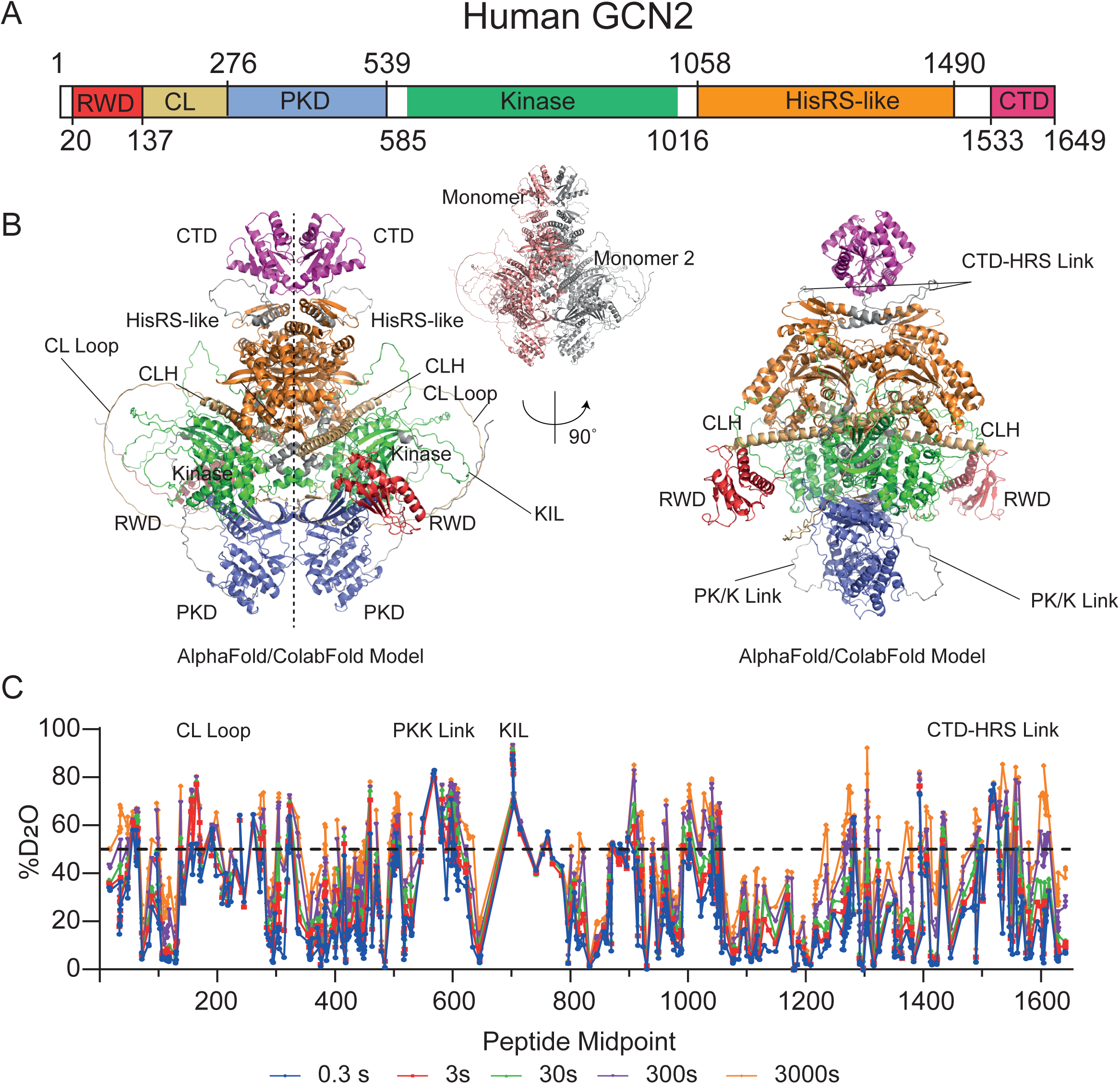
Predicted Model of GCN2 generated by AlphaFold/Collabfold and comparison to HDX-MS data. **(A)** Domain organisation and amino acid residue boundaries for human GCN2. CL = Charged Linker, PKD = pseudokinase domain, HisRS = Histidyl synthetase like domain, CTD = C-terminal Domain **(B)** Views of the GCN2 dimer generated using AlphaFold/Colabfold. CLH = charged linker helix, KIL = kinase insertion loop. **(C)** HDX-MS data for APO GCN2. Typically, regions where deuterium uptake is greater than 50% (shown as the dashed line) at 0.3s incorporation are likely unstructured loops with his flexibility.

**Table 1.0.** HDX-MS Statistics.

|  | Apo | GDC | Dab | Enc | GCN2iB |
| --- | --- | --- | --- | --- | --- |
| # of Peptides | 687 |  |  |  |  |
| Coverage | 95.4% |  |  |  |  |
| Av. Length | 13.2 a.a. |  |  |  |  |
| Redundancy | 5.5 |  |  |  |  |
| Time Points | 0.3s*/3s/30s/300s/3000s |  |  |  |  |
| Replicates | 3 per time point. 5 ND. |  |  |  |  |
| Controls | Non-Deuterated and Maximally Deuterated Control |  |  |  |  |
| Replicability | Average % STDEV for technical triplicate = 1.00% |  |  |  |  |
| Significance threshold | >5σ & t test <0.05 | | | | |

\* 0.3s timepoint created by conducting a 3 s timepoint on ice.

Using HDX-MS, we found that Dabrafenib, Encorafenib and GCN2iB all produced distinct changes in solvent exchange, and that GDC-0879 caused no significant changes in exchange rates (See Figures 3 and 4). Dabrafenib, Encorafenib and GCN2iB all bound within the kinase of domain of GCN2, causing large reductions in solvent exchange in peptides surrounding the ATP-binding pocket (residues 587-592, 803-818, 836-876, 902-914) (see Fig 3A, B, C and Fig 4). Surprisingly, compound binding was accompanied by several additional exposures within the kinase domain (861-871, the αC helix (Fig 3C)). Distal from the kinase domain, there are further exposures in GCN2 Unique N-terminal Subdomain of the HisRS (1010-1056) (Figure 3E), the HisRS Domain (1369-1390) (Figure 3E), and the CTD (1591-1633) (see Fig 3D). Comparing the patterns of changed solvent exchange for each of the compounds, Dabrafenib and Encorafenib were most similar in their patterns in exchange, with GCN2iB recreating many of these features also.

**Figure 3:**
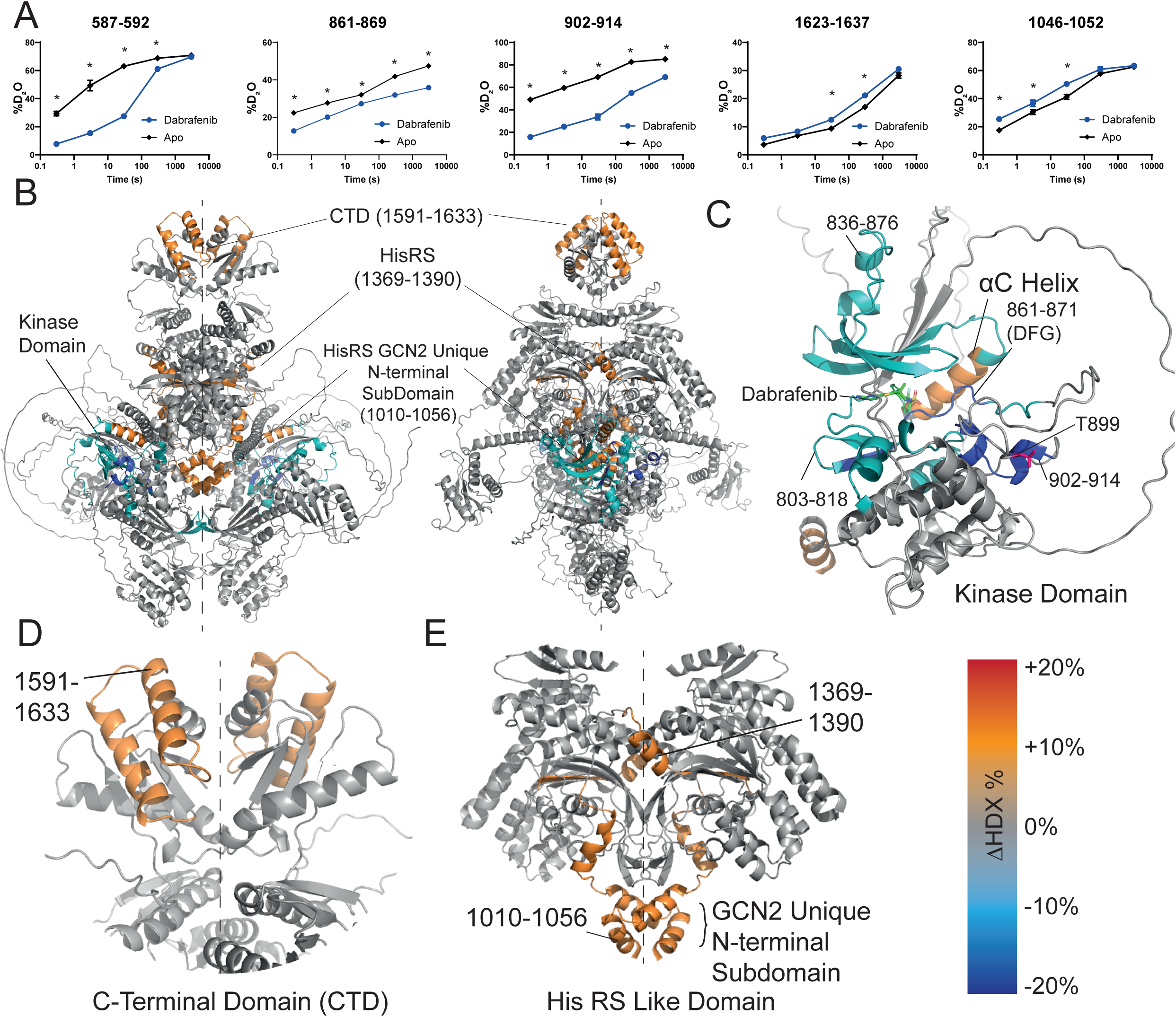
HDX-MS changes associated with Dabrafenib binding. **(A)** Selected peptides showing the effect of Dabrafenib binding to GCN2. * = Greater than 5% & 0.5Da change, passes t-test at p=0.05. Error bars may be smaller than the points on the graph. **(B)** Dabrafenib binding mapped onto the entire GCN2 model. Cooler colours represent areas of ‘protection’ (i.e., slower solvent exchange rate), warmer colours show ‘exposure’ (i.e., increased solvent exchange rate). **(C)** Inset of the kinase domain of GCN2 showing HDX changes on Dabrafenib binding. T899, a “bell weather” site of autophosphorylation and activation is highlighted. **(D)** The C-Terminal Domain of GCN2, highlighting the exposure of the helical hairpin found in residues 1591-1633. **(E)** The His-RS Like Domain with the GCN2 Unique N-terminal Subdomain included, highlight the exposure occurring in this domain. Dashed lines represent 2-fold symmetry of the dimer.

**Figure 4:**
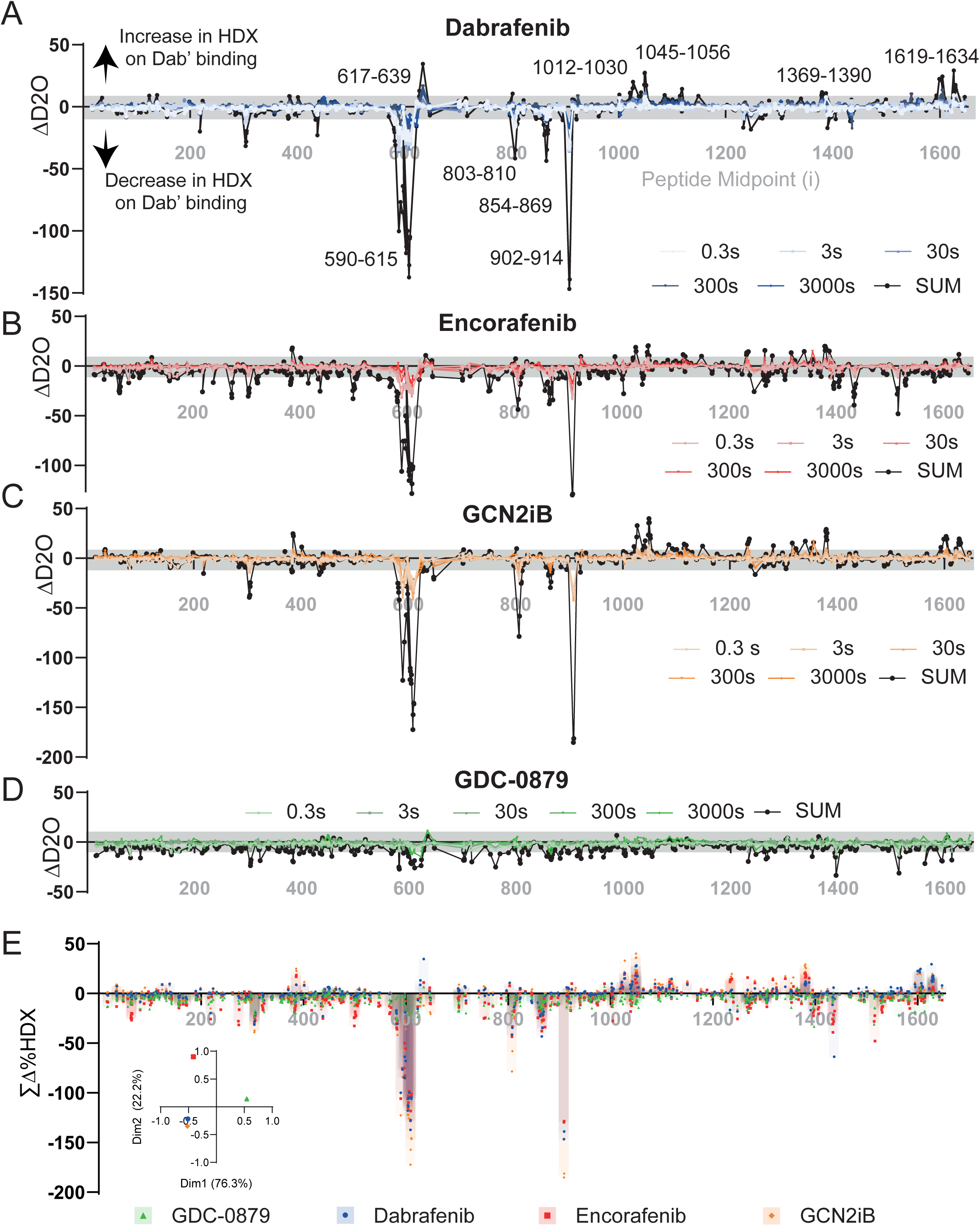
Comparison of HDX-MS changes with (A) Dabrafenib (B) Encorafenib (C) GCN2iB (D) GDC0876. Each point in the midpoint of a peptide with the five-points in gradually more saturated colours. Black points/line are the SUM difference across all timepoints. **(E)** Amalgamated changes mapped onto one another. Inset is a principal component analysis comparing all the data summed data.

### GCN2 can engage in “para-autophosphorylation”

Current models of GCN2 activation assume that GCN2 can only engage in “cis-autophosphorylation” whereby a GCN2 kinase domain can phosphorylate its ‘own’ Thr899 residue (i.e. autophosphorylation within the same polypeptide) or perhaps (with sufficient domain rearrangement) “trans-autophosphorylation” with the Thr899 residue of the dimer partner. We wished to explore whether an activated GCN2 dimer was capable of phosphorylating inactive GCN2 dimers, as this would provide an alternative explanation for how inhibitor-bound GCN2 dimers were capable of becoming phosphorylated. First, we recombinantly expressed and purified a kinase dead mutant of GCN2 – GCN2^K619R^ (See Supplementary Figure 2) ^34^. We found the mutant to be a stable dimer (see Supplementary Figure 2), with a melting temperature and profile very similar to that of wtGCN2 (wt GCN2 47.4 ±0.1 °C, GCN2^K619R^ 47.3 ±0.1 °C (see Supplementary Fig 2C)) and no detectable autophosphorylation (even after 2 h incubation) or substrate (full length eIF2α) activity, and negligible ATPase activity (See Figure 5A, 5B). Next, to see whether wtGCN2 could phosphorylate GCN2K619R, we immobilised N-terminally strep-tagged wtGCN2 on beads and then incubated the immobilised wtGCN2 with an excess of GCN2 K619R and ATP. Surprisingly, we were able to detect pT899 on catalytically inactivated GCN2 K619R. Furthermore, when we conducted a concentration gradient of Encorafenib in this system, we were able to observe a similar concentration dependent increase in pT899 on GCN2 K619R (See Figure 5D, 5E).

**Figure 5:**
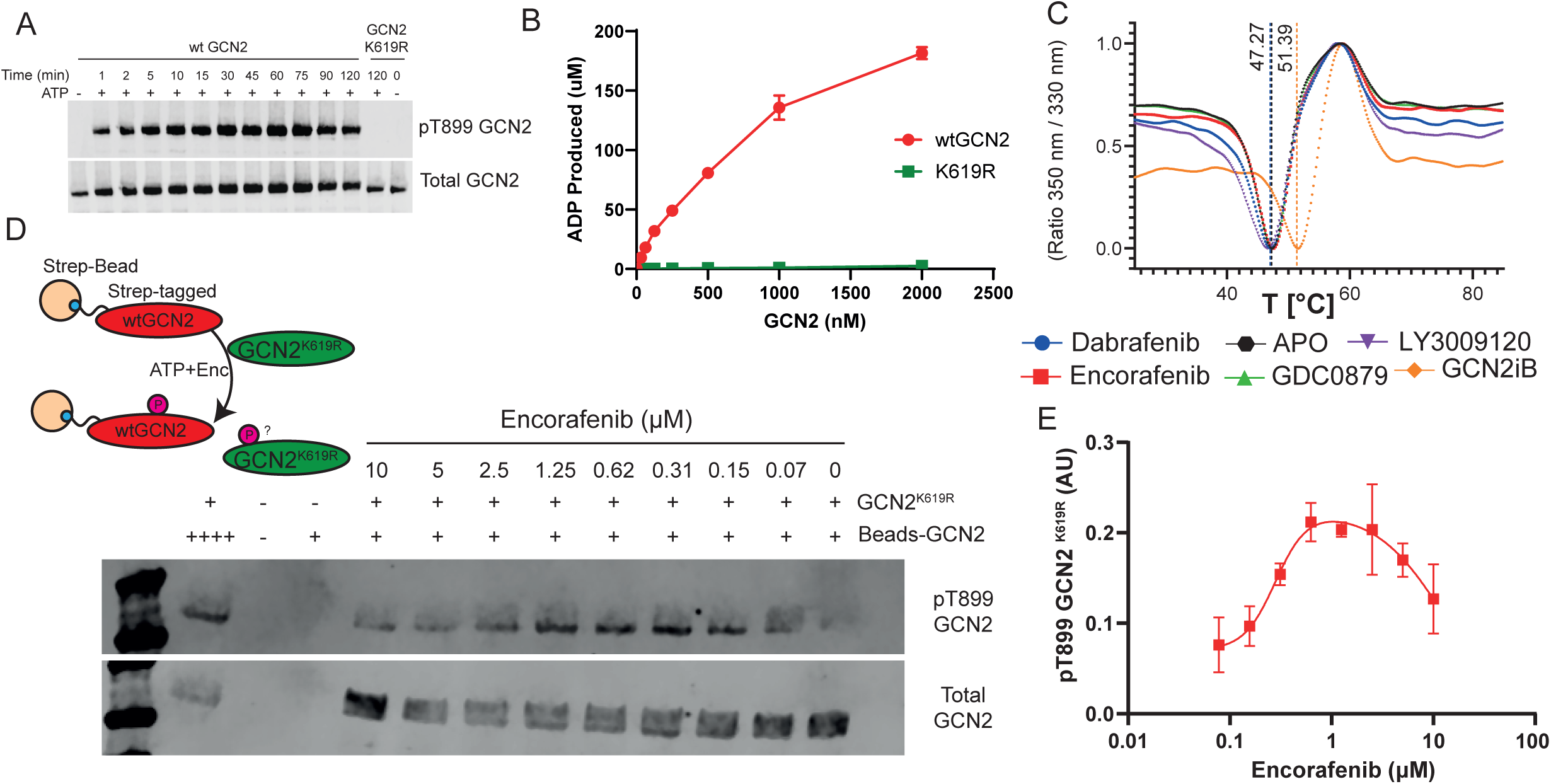
GCN2 is capable of phosphorylating in *trans*. **(A)** Autophosphorylation activity comparison of wtGCN2 with GCN2^K619R^. **(B)** Comparison of ATPase activity as measured using ADPGLO between GCN2 and GCN2^K619R^. **(C)** Thermal Unfolding Assay for GCN2^K619R^ on compound binding. **(D)** Western Blot analysis of trans-phosphorylation of GCN2. **(E)** Compound-driven trans phosphorylation of GCN2^K619R^ by immobilised wtGCN2 by Encorafenib.

## Discussion

In this manuscript we use biochemical and structural data to present a model of GCN2 activation by small molecule ATP-competitive inhibitors of RAF and GCN2iB. Our recent studies in collaboration with Gilley *et al.* () show that small molecule BRAF inhibitors were effective GCN2 activators in cells, and this was confirmed using our *in vitro* kinase assay. Here we illustrate how these compounds may facilitate GCN2 activation via a mechanism of allosteric activation which is propagated through the enzyme.

We found that there was a disconnect between the concentration of compound where a peak of GCN2 autophosphorylation was observed and eIF2α phosphorylation could be found. This observation appeared inconsistent with the previous models proposed, where compound occupancy of a single GCN2 monomer facilitated activation of the other monomer within a single GCN2 dimer, thus allowing GCN2 to phosphorylate its substrate i.e., trans-activation, as has been proposed for BRAF/MAPK pathway kinases ^35,36^. If this previously suggested model were correct, there should be a typical initial activation autophosphorylation event followed by multiple substrate phosphorylation reactions, as the compound would be acting as an allosteric activator, producing 1:1 correlated p-eIF2α/p-GCN2 levels– however this was not observed. Furthermore, it required a complex array of allosteric interactions that facilitated occupancy of only a single kinase domain, whilst allowing the second kinase domain to remain free of compound. This disagreed with many studies which detailed that GCN2 requires multiple intramolecular rearrangements to achieve autophosphorylation ^32,33,37,38^, and that there is a reorientation of the kinase domains from an inactive antiparallel to active parallel orientation during activation^39^. One explanation for this discrepancy may be that experiments exploring paradoxical activation of GCN2iB were conducted with isolated fragments of GCN2 where the complex networks of intramolecular allostery that govern GCN2 activation by *e.g.,* tRNA, would presumably be absent ^23,24^. Given this apparent discrepancy, we investigated how compounds activate GCN2 using structural mass spectrometry.

Using HDX-MS, we were able to observe where these compounds were binding to full-length human GCN2 and determine what wider allosteric effects may result from their binding. HDX-MS has been used in several studies to determine the mode of interaction of kinase inhibitors ^40–43^, and previous investigations using HDX-MS and GCN2 had revealed some of the likely structural changes which occur in GCN2 upon activation via P-stalk binding ^18^. We saw a similar pattern of structural changes upon compound binding of Encorafenib, Dabrafenib and GCN2iB with a high degree of solvent exchange reduction within the kinase domain, indicative of compound binding.

To note, GCN2 has three potential ATP-binding sites: namely, the canonical kinase domain, and the more cryptic sites in the pseudokinase and HisRS domains. We only observed reductions in solvent exchange (indicative of a direct interaction) in the kinase domain site; this is consistent with our previous modelling predictions and expression of gatekeeper mutations in cells (Gilley et al.) However, there were also several unexpected increases in solvent exchange distal to the ATP-binding site of the kinase domain, and thus these increases in HDX are likely to be the result of the breaking of auto-inhibitory hydrogen bonding networks – similar events have been observed before with other allosteric activation events ^44^.

Previous mutagenesis studies of GCN2/Gcn2 have provided insight into the network of interactions which facilitate autoinhibition and subsequent activation of the enzyme ^30,38,45,46^. These hydrophobic residues and autoinhibitory interactions span the enzyme but form a nexus on L869 (L856 in yeast Gcn2) – a residue that follows directly after the ‘DFG’ of the activation loop of the kinase domain, and strands β3 or β5 and the helix αC ^30^. Our HDX data shows all the activating compounds influencing these elements of the kinase domain, and thus it may be triggering this network of interactions. Most apparent for its disruption of the αC helix was Dabrafenib, binding of which resulted in an increase in solvent exchange in this area. Dabrafenib binding may disrupt the hydrophobic node formed by I633, V637 and L639, and may also influence by the breaking of the inactivating salt bridge which forms between E636 (also in the αC and R847 (in the catalytic loop)^46^ – all these regions saw changes in the solvent exchange upon Dabrafenib, Encorafenib and GCN2iB binding.

Other regions crucial for GCN2 activation are the C-terminal domain (CTD) and the histidyl-tRNA synthetase (HisRS) domains - the interplay between these two domains has been demonstrated to be key for GCN2 activity ^37^. Within the HisRS domain we saw increases in solvent exchange in residues 1369-1390 (Fig 3E) – this region was found be the most protected in a recent publication mapping tRNA biding to the isolated HisRS domain ^33^ – thus the H-bonds network appears to be melting the structure in this region typically associated with tRNA interaction. This may be indicative of a “priming” of the site for tRNA binding. Mutagenesis of D1353 (yeast D1327) to either alanine or lysine completely abrogates GCN2 activity ^37^ –the recently published human HisRS domain structure ^33^ shows D1353 near to the region of exposure produced upon RAFi/GCN2iB binding, which may suggest that this region acts a gatekeeper for activation of the enzyme.

In our AlphaFold/ColabFold model of GCN2, the N-terminus of the HisRS domain sits in very close proximity to both the PKD and KD domains – this may provide insight into how binding within the kinase domain is propagated through the HisRS domain. The HDX-MS data shows that there are increases in solvent exchange on compound binding in a helical bundle found within the first 50 residues of the HisRS domain (1010-1055), and proximate to this bundle is a cradle of beta-hairpin loops of the PKD β3/β4 strands (residues 299-309). The helical bundle – referred to as the “GCN2 Unique N-terminal Subdomain” by Yin *et al.* ^33^, sits directly above the ATP binding loop of the HisRS domain -and in our model, may facilitate the propagation of allostery. Given these compounds appear to be “priming” GCN2 for activation by either the CTD or HisRS domains, investigating whether these routes to activation are promoted by compound binding requires further investigation.

Dabrafenib, Encorafenib and GCN2iB binding in the kinase domain caused structural changes in the CTD, even though it is approximately 80 Å away from the binding site with our model. This increase in HDX suggests a breakdown of the hydrogen bonding in this region – although interpreting this change in solvent exchange is difficult. Mapping the peptides covering the region of the (1591-1637) to the murine and yeast crystal structures produced of the CTD dimer (Supplementary Figure 3) ^47^, we see that the exposure is largely confined to a pair of helices that present away from the dimeric interface. A key residue for maintaining the CTD dimer, L1587, is nearby, but does not show increased exposure on compound binding. Truncation analysis in yeast found that part of the CTD could directly bind to the ribosome ^48^ – although the construct used did not contain the 1591-1637 helix – its exact function is currently unknown. Overall, our HDX-MS data supports a model of catalytic inhibition coinciding with a partially activated structure where many known points of regulation have become more flexible.

The ‘activation’ of GCN2 has frequently been synonymous with both autophosphorylation of T899 and eIF2α phosphorylation - however, our analysis has shown that phosphorylation can effectively be de-coupled from eIF2α by these ATP-competitive compounds. Furthermore, “true” activation of GCN2 by tRNA or ribosomal binding has often been associated with large rearrangements of GCN2, especially with the kinase domain adopting an inactive antiparallel stance which then rearranges to an active parallel dimer via activation^39^. Our HDX-MS data show that these compounds activate GCN2 but do not appear to induce the large-scale rearrangements that would be necessary to facilitate a shift from antiparallel to parallel arrangement of the kinase domain – strengthening the possibility that these compounds are essentially catalytically inhibiting, while facilitating phosphorylation of GCN2.

Finally, we observed that GCN2 may be capable of phosphorylating in *trans* – and that the ATP-competitive compounds can make GCN2 a better substrate for *trans* phosphorylation by presenting a pseudo-active state. However, further research will be required to determine whether this sequence of events is what occurs in a cellular context. It raises the possibility that there is the potential for a positive feedback loop occurring whereby activation of minor populations of GCN2 allows for a burst of activating trans phosphorylation events and the rapid amplification of the ISR – something which may be required if the initiating event – frustrated ribosomal collision^1522^, is a relatively low-frequency/density occurrence in the cell.

The observation that a wide array of ATP-mimetic kinase inhibitors, including those that are clinically approved and are thought to act through alternate signalling pathways can activate GCN2 within a cellular context has implications for drug discovery, not least in the efforts to target GCN2. This is especially noteworthy in cancer cells where, in response to the compounds, there will be a selective pressure to alter GCN2 concentrations to facilitate cell growth^49,50^. If it is beneficial within certain pathologies to inhibit GCN2, it may be worthwhile employing a drug discovery programme that avoids ATP-competitive compounds and instead attempts to inhibit GCN2 by allosteric means, or targeted protein degradation (i.e., proteolysis targeting chimeras (PROTACs)).

---

**Supplementary Figure 1:**
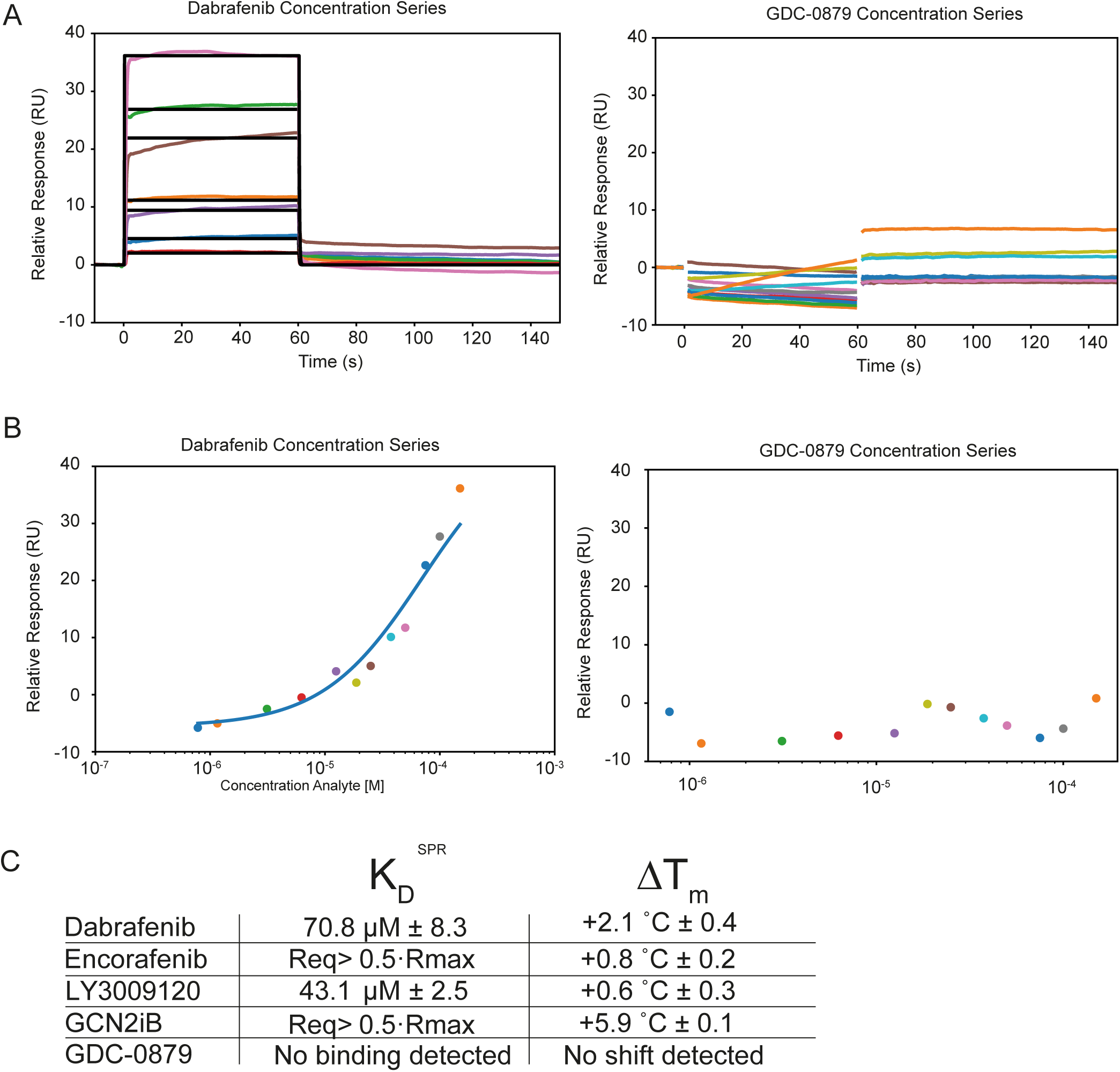
Binding of Compounds to GCN2. **(A)** SPR traces of binder Dabrafenib and non-binder GDC-0879 **(B)** Steady-State analysis of compound binding derived from SPR **(C)** Amalgamated compound characteristics. *Analytes including Encorafenib and GCN2iB demonstrated binding with SPR, producing a concentration dependent response, but at the analytes at higher concentrations shows superstoichiometry or binding without complete saturation of the ligand in the concentration range used ^51^. The affinities, as determined by kinetic analysis, could not be reliably determined.

**Supplementary Figure 2:**
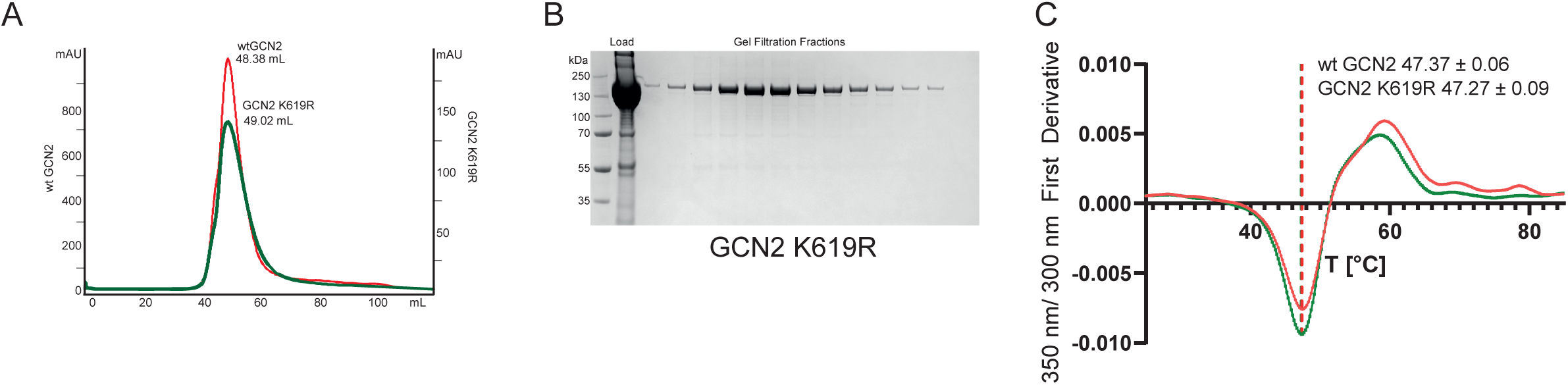
Characterisation of GCN2^K619R^. **(A)** Gel Filtration profile of wtGCN2 (red) and GCN2^K619R^ (green) showing similar elution profiles **(B)** SDS-PAGE of Gel Filtration of fractions of GCN2^K619R^ **(C)** Thermal denaturation profile of wtGCN2 (red) and GCN2^K619R^ (green).

**Supplementary Figure 3:**
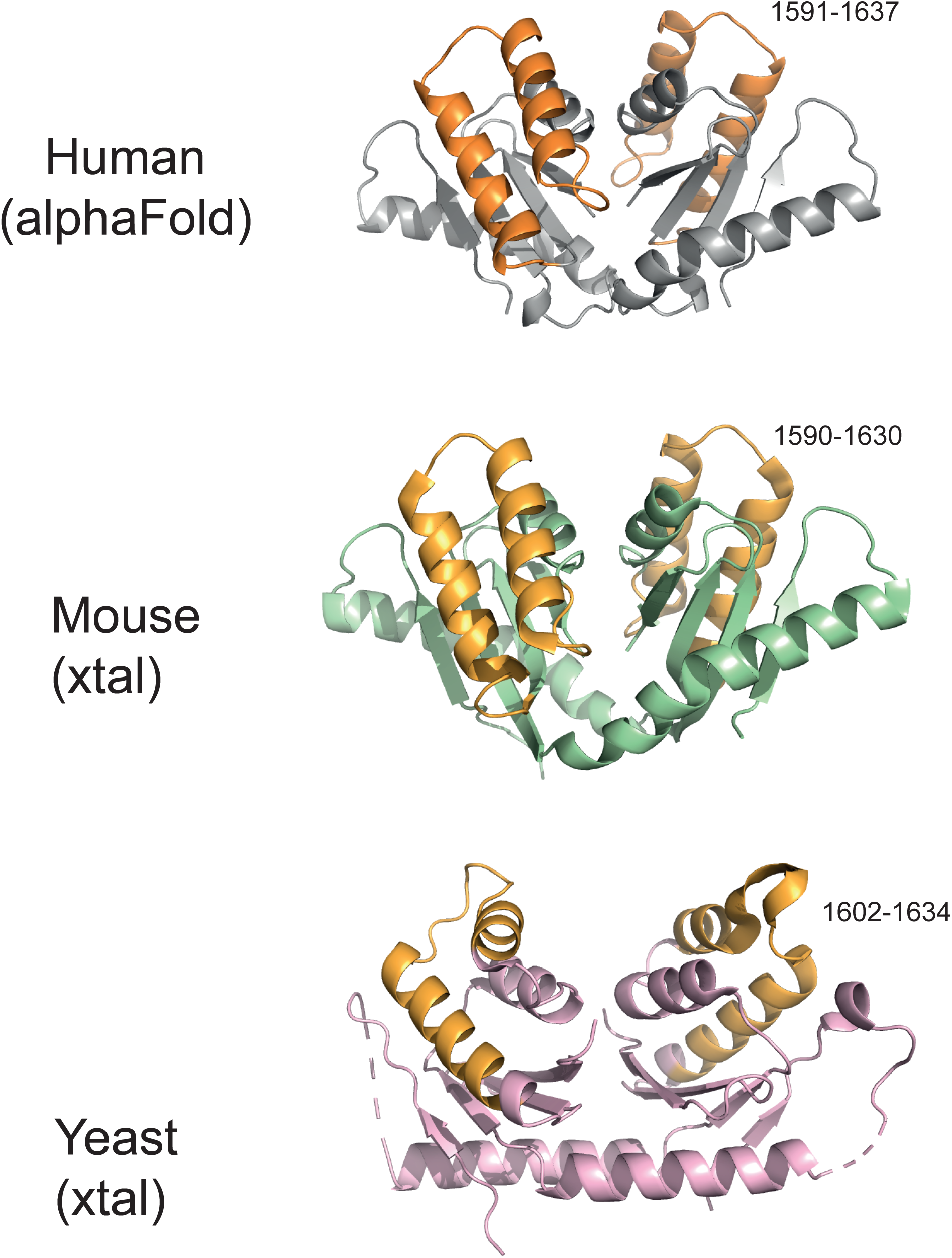
Comparison of CTDs. The CTDs of Human (produced by Alphafold), Mouse (crystal structure PDB: 4OTN), and Yeast (4OTM), with the helix-turn-helix motif which exhibited an increase in HDX-MS on compound binding shown in orange.

## Acknowledgements

GRM and GN are supported by a Baxter Fellowship from the University of Dundee, and a TENOVUS grant T21/01 119615, and Biotechnology and Biological Sciences Research Council (UKRI-BBSRC) Capital Equipment Grant BB/V019635/1. VV supported by MRC iCase Studentship (MR/R01579/1). R.G. and S.C. were Plexxikon Inc and Institute Strategic Programme Grants BB/J004456/1, BB/P013384/1 and BB/Y006925/1 from the UKRI-BBSRC.

## Competing Interests Statement

The authors announce no competing interests. The initial stages of this work (i.e. identification of RAFi/GCN2 activation in a cellular context) were supported by sponsored research collaboration in SJC’s laboratory funded by Plexxikon Inc.

## Materials and Methods

### Antibodies and Reagents

Primary Anitbodies: Total EIF2S1 (eIF2α) (1:1000 WB, SC133132, Santa Cruz), phosphoserine 51 EIF2S1 (1:500 WB, 9721S, Cell Signalling Technology), Total GCN2 (1:1000 WB, MA5-32704, Invtirogen), phosphothreonine 899 GCN2 (1:1000 WB, AB75836, Abcam) 6X His Tag (1:1000 WB, R931-25, Invitrogen),

Secondary Antibodies: IRDye® 680RD Goat anti-Mouse IgG Secondary Antibody (926-68070, Licor), IRDye® 800CW Goat anti-Mouse IgG Secondary Antibody (926-32210, Licor), IRDye® 800CW Donkey anti-Rabbit IgG Secondary Antibody (926-32213, Licor), IRDye® 680RD Goat anti-Rabbit IgG Secondary Antibody (926-68071, Licor).

### GCN2 Expression and Purification

Expression and purification of recombinant human GCN2 was conducted as described previously^18^. Briefly, human GCN2 cloned (uniprot ID: Q9P2K8) was cloned into a baculoviral vector with an N-terminal twin StrepII tag followed by a TEV protease site. Plasmids were then integrated into the EMBacY baculoviral DNA, and the baculovirus amplified over three rounds. GCN2 was then expressed in *Sf9* cells by infecting cells at a density of 1.5x 10^6^ cells/mL with 15 mL of virus being used per 500 mL of cells. Cells were grown at 27 °C for 55 hours and were then harvested via centrifugation, washed in ice cold PBS and snap frozen in liquid nitrogen. Pellets were then thawed on ice with 100 mL of Lysis Buffer A (20 mM Tris pH 8.0, 150 mM NaCl, 5% v/v glycerol, 2 mM β-mercaptoethanol (BME), and one cOmplete EDTA-free protease inhibitor tablet per 50 mL of Buffer). Cells were then lysed via probe sonication, on ice, for 5 min (10 s on/ 10 s off), and then Benzonase at 2 U/mL was added. Lysate was then centrifuged at 140,000 *g* at 4 °C for 45 minutes. Proteins were purified using an AKTA protein purification system (Cytiva). The supernatant was then filtered through a 0.2 µm syringe filter before being loaded onto 2 x 5 mL StrepTrap HP Columns (GE Healthcare Life Sciences 28-9075-47), equilibrated in Step A Buffer (20 mM Tris pH 8.0, 150 mM NaCl, 5% v/v glycerol, 2 mM BME) at a flow rate of 4 mL/min. Protein was then eluted through a gradient of Strep B Buffer (20 mM Tris pH 8.0, 150 mM NaCl, 5% v/v glycerol, 2 mM BME, 6 mM desthiobiotin). Peak fractions were analysed using SDS-PAGE and assessed for relative purity. GCN2-containing fractions were then diluted using Q_0_ Buffer (20 mM Tris pH 8.0, 5% v/v glycerol) to a NaCl concentration of ∼50 mM before being loaded onto a 5 mL HiTrap Q HP column (Cytiva 17115401), equilibrated in QA Buffer (20 mM Tris pH 8.0, 50 mM NaCl, 5 & v/v glycerol, 2 mM BME), at a flow rate of 4 mL/min. The column was then washed with 100 mL QA Buffer, before a gradient of QB Buffer (20 mM Tris pH 8.0, 1 M NaCl, 5% v/v glycerol, 2 mM BME) was applied. GCN2 containing fractions were then pooled and concentrated using a 50 mL Amicon Centrifugation Concentrator (50k MWCO) to a volume of ∼1 mL before being injected onto a Hi Load 16/60 Superdex 200 column (Cytiva/GE Healthcare 28989335), equilibrated with GF buffer (20 mM HEPES pH 7.5, 150 mM NaCl, 2 mM tris(2-carboxyethyl)phosphine (TCEP)), at a flow rate of 1 mL/min. GCN2 containing fractions were concentrated using a 50 mL concentrator to a concentration of ∼3 mg/ml before being aliquoted and snap frozen in liquid nitrogen.

### GCN2^K619R^ Expression and Purification

Human wtGCN2 was mutated via QuickChange mutagenesis to K619R (Forward Primer: GGACGGTTGCTGCTACGCTGTGAGGCGTATCCCCATCAACCCTGC, reverse: CGGGAAGCAGGGTTGATGGGGATACGCCTCACAGCGTAGCAGC). Expression and purification proceeded as per wtGCN2, except after the Strep-Tag-purification, GCN2 K619R was cleaved for 16h at 4°C with TEV protease to remove an N-terminal His-Strep Tag. Purification then proceeded as per wtGCN2 (Q Column and Gel Filtration in the same buffers). Fractions were concentrated to 5.5 mg/mL and stored as per wtGCN2.

### eIF2α Expression and Purification

Expression and purification of recombinant human eIF2α was conducted as described previously^18^. DNA encoding full-length human eIF2α (NCBI reference number: NP_004085.1) was inserted into the vector pOPTH with an N-terminal His6 tag followed by a TEV protease site. The plasmid was transformed into chemically competent BL21 Star (DE3) cells, and cells were grown overnight before being inoculated to a 50 mL starter culture in 2xTY media containing 0.1 mg/mL Ampicillin. The starter culture was incubated at 37 °C for 90 minutes, then 10 mL starter culture was added to 4 x 900 mL 2xTY media containing Ampicillin. Cultures were incubated at 37 °C until the optical density reached 0.7, and then protein expression was induced by the addition of 0.3 mM isopropyl β-D-1-thiogalactopyranoside (IPTG). Cells were grown for a further 3 hours at 37 °C before being harvested, washed with ice-cold phosphate-buffered saline and frozen in liquid nitrogen. Bacterial cell pellets were lysed in 100 mL Lysis Buffer (20 mM Tris-HCl pH 8.0, 100 mM NaCl, 5 % v/v glycerol, 2 mM BME 0.5 mg/mL Lysozyme (Sigma L6876), 2 U/mL Benzonase, one cOmplete EDTA-free protease inhibitor tablet (Roche 04693132001) per 50 mL of Buffer). Cells were lysed using a probe sonicator for 5 minutes (10 s on/ 10 s off) and then centrifuged at 140,000 *g* for 45 min at 4 °C. The supernatant was filtered through a 0.2 µm syringe filter before being loaded onto a 5 mL HisTrap HP Column (Cytiva 17524801) equilibrated in Ni A Buffer (20 mM Tris pH 8.0, 100 mM NaCl, 5 % v/v glycerol, 10 mM imidazole pH 8.0, 2 mM BME), followed by the elution of protein via a gradient of Ni B Buffer (20 mM Tris pH 8.0, 100 mM NaCl, 5% v/v glycerol, 200 mM Imidazole pH 8.0, 2 mM BME). Protein purification then proceeded as described for GCN2. Proteins were concentrated to ∼10 mg/mL and then snap frozen in liquid nitrogen.

### Thermal Unfolding Assays

Thermal Unfolding Assays were conducted using a NanoTemper Panta using Prometheus Standard Capillaries (NanoTemper) (PR-C002). Briefly, GCN2/GCN2 K619R was defrosted, and centrifuged at 20,000 *g* for 10 minutes, and then diluted to 0.3 mg/mL in 20 mM HEPES pH 7.5, 150 mM NaCl, 1 mM TCEP Buffer was incubated with compounds on ice for 45 minutes before being analysed. A gradient of 1 °C/min was applied to the sample starting from 25 °C to 85 °C. Fluorescence at 330 nm and 350 nm was monitored using an excitation wavelength of 280 nm. Data analysis and identification of inflection points/TMs was conducted using the Panta Analysis Software (NanoTemper).

### Surface Plasmon Resonance (SPR) Studies

SPR studies were carried out using Biacore T-200 (GE Healthcare) and Biacore 8K+ (Cytiva) using Series S NTA Sensor Chips (Cytiva). The running buffer was HBS-P+ (Cytiva) with 1% DMSO, with the flow cell temperature held at 25 °C, and the sample compartment temperature held at 10 °C. His-tagged GCN2/GCN2^K619R^ at a concentration of 0.05 mg/mL was immobilised using custom capture-immobilization method to an RU of ∼8600RU. Ligands were captured on pre-conditioned chips activated in 2 steps first by NiCl2 followed by a 1:1 mixture of 0.4M 1-ethyl-3-(3-dimethylaminopropyl)-carbodiimide hydrochloride (EDC) and 0.1M N-hydroxysuccinimide (NHS) for 7 min at a flow rate of 10 μL/min.

Multi-cycle analyte binding studies was performed using 12-point dose response curve for each analyte. Two concentration series starting from 150 µM were prepared for each analyte from 10 mM stocks in 100% DMSO. Dilutions of analyte were prepared in 1xHBS-P+ at final concentration of 1%-1.5% DMSO. Analysis step included and association time of 60s and dissociation time of 120 s, at a flow rate of 40 μL/min. Data was collected at a rate of 10 Hz using T200 Control software Version 2.0.1 in High performance mode and analysed using Biacore Insight Evaluation Software™-Version 5.0.18.22102. Solvent correction was performed using 6-point linear concentration range between 0.2%-1.8% DMSO. Data analysis was performed on the solvent corrected, blanked curves using a 1:1 binding (Rmax constant at Ymax) model and steady state affinity (constant Rmax, 10RU per 100Da) model for a response window of 5 s at 5 s before end of injection (equilibrium analysis) for relevant concentrations. Steady state affinities and kinetic data have been used for qualitative assessment of target binding and the rank ordering of relative affinities.

### GCN2 Activity Assays

#### ADP-GLO

Using a PROMEGA ADP-GLO Kinase assay, 100 nM GCN2 was incubated with 50 µM polyarginine-serine repeat peptide (Sigma-Aldrich SRP0683) or 10 µM eIF2α with 500 µM ATP in 1 x Kinase Reaction Buffer (20 mM Tris pH 8.5, 0.2 mM EDTA, 0.1 % Tween 20, 100 mM NaCl, 0.001% Brij-35, 0.5% glycerol, 10 mM MgCl2, 0.1 mg/mL BSA) for 1 hour at room temperature, until the reaction was quenched using the ADP-GLO Reagent.

#### Phos-tag Gels

Phos-tag gel assays were also conducted to facilitate relative substrate phosphorylation and autophosphorylation levels. 400 nM GCN2 was incubated with 14 µM eIF2α and 250 µM ATP and 50 µM deacylated tRNA in 20 mM Tris pH 8.0, 150 mM NaCl, 10 mM MgCl2 at 30 °C. Reactions were quenched through mixing with 2xSDS-PAGE Loading buffer with 1 mM ZnCl2. 10 µL of reactions were loaded onto SuperSep Phos-tag (50 µmol/L) 100 x 100 x 6.6 mm (17 well) gels (Fujifilm). Gels were run in the manufacturer’s suggested 100 mM Tris pH 7.8/ 100 mM MOPS/ 0.1% SDS / 5 mM Sodium Bisulfite solution. Gels were held at a constant 100 V for 2 h prior to staining with InstantBlue Coomassie Stain (AbCam).

#### Western-Blot

10 nM GCN2 was incubated at 30 °C for 20 min with 2 µM eIF2α and 10 µM ATP in varying deacylated tRNA concentrations. Samples were then quenched via the addition of SDS-PAGE loading buffer and heating to 95 °C for 3 min. Samples were then run in a MES buffer SDS-PAGE experiment using a 15 well 5-12% Bis-Tris Gel, run at 180 V for 35 min at RT. Gels were then briefly incubated in 20% ethanol solution prior to a transfer to a nitrocellulose membrane using the iBlot 2 Transfer system. Membranes were then blocked using 5% Milk TBST solution for 1 hour at RT, before being washed three times 10 min incubation with TBST. Primary antibodies were diluted as follows in 4% BSA TBST: total eIF2alpha (Santa Cruz Biotechnology sc-133132) 1:1000, phospho-eIF2S1 Ser51 (Cell Signalling Technology 9721) 1:500, total GCN2 (Invitrogen JA03-83) 1:1000, pThr899 GCN2 (Abcam AB75836) 1:1000. Primary antibody solutions were incubated with membranes in heat-sealed bags with gentle agitation for 1 h at RT or overnight at 4 °C. Membranes were then washed 3 x 10 min with TBST. Secondary antibodies (LICOR IRDye 800 CW (926-32210 and 926-32213) were then prepared to manufacturer’s instructions (0.5 µL in 5 mL 4% BSA TBST) and incubated for 1 h in a 50 mL Falcon tube on a roller. Membranes were washed with 3 x 10 min TBST before imaging with a LICOR Odyssey FC. Images were captured and quantified in Image Studio Ver 5.2.

#### Immobilised GCN2^K619R^ Activity Assay

25 µL (per reaction) of IBA Strep-Tactin Sepharose resin was washed with Reaction Buffer (20 mM Tris pH 7.5, 150 mM NaCl, 50 mM MgCl_2_, 1 mM TCEP) five times via resuspension and centrifugation. 100 µL of 20 nM Strep-Tagged wtGCN2 was immobilized on resin for 1 h at 4 °C with agitation, followed by five more washes with twice the bead volume. Resin was then incubated with 2 µM non-tagged GCN2^K619R^, 10 µM and compound in a 50 µL final volume (i.e., 50% beads with 20 nM wt-GCN2 bound for a final concentration of 10 nM wt-GCN2). The reaction was incubated for 20 min at 30 °C. The reaction was quenched via separating the beads and substrate GCN2^K619R^ via centrifugation using a Cytiva MicroSpin Column, and the flowthrough was then mixed with SDS-PAGE loading buffer and boiled at 94 °C for 3 minutes. The beads were then further washed twice to remove residual GCN2^K619R^, followed by elution using Elution Buffer (20 mM Tris pH 8.0, 150 mM NaCl, 5 mM desothiobiotin, 1 mM TCEP). GCN2 phosphorylation was measured using Western Blots as described above.

### Hydrogen Deuterium Exchange Mass Spectrometry

#### Sample Preparation

5 µM GCN2 was incubated with or without 100 µM Compound (as stated) for 30 minutes in Protein Dilution Buffer (20 mM HEPES pH 7.5, 150 mM NaCl, 15 mM MgCl2, 2 mM TCEP). 5 µL of this sample was diluted with 45 µL of Deuteration Buffer (20 mM HEPES pH 7.5, 150 mM NaCl, 15 mM MgCl2, 2 mM TCEP, 1% DMSO, 96.5 % D2O) (final D2O concentration = 85.95%) for timepoints 3/ 30/ 300/ 3000 s, before being quenched with 20 µL of ice cold Quench Solution (6 M Urea, 2% Formic Acid), and being snap frozen in liquid nitrogen and stored at – 70 °C. A further timepoint, 0.3 s, was achieved by incubating both the sample and deuteration buffer on ice and then conducting a 3 s exchange reaction. Each exchange reaction was conducted independently four times.

#### Data Acquisition

Data acquisition was conducted broadly as previously described^44^. Samples were rapidly thawed at room temperature and then injected into automated HDX-MS fluidics and UPLC manager system (Waters). Samples were loaded into a 50 µL loop and then subsequently digested using a Waters Enzymate BEH Pepsin Column (Part No. 186007233) in a 0.1 % Formic Acid solution with a flow rate of 200 µL/min at 20 °C. Peptic peptides then flowed onto a Waters ACQUITY UPLC BEH C18 VanGuard Pre-Column at 1 °C (Part No. 186003975). After digestion, the flowpath was changed to elute the peptides via a Waters ACQUITY UPLC BEH C18 1.7 µm 1.0 x 100 mm reverse phase column (Part No. 186002346). A gradient from 0-85% 0.1% Formic Acid/ Acetonitrile, conducted at 1 °C with a 40 µL/min flowrate was used to elute deuterated peptides which were then ionised using an ESI source. Mass spectrometry data were collected using a Waters Select Series cIMS instrument, from a 50-2,000 m/z range with the instrument in HDMSe mode. A single pass of the cyclic ion mobility separator (with a cycle time of 47 ms) was conducted. A blank sample of protein dilution buffer with quench was run between samples, and carryover was routinely checked to be <1% intensity of the prior sample.

#### Data Analysis

Peptide sequence identification was conducted using Protein Lynx Global Server (Waters). Minimum inclusion criteria were a minimum intensity of 5000 counts, minimum sequence length 5, maximum sequence length 35, a minimum of 3 fragment ions, a minimum of 0.1 products per amino acid, a minimum score of 5.0, a maximum MH+ Error of 10 ppm. Subsequent analysis and determination of deuteration values conducted using HDExaminer (Sierra Analytics/Trajan).

## CRediT Author Contribution

Graham Neill: Conceptualisation, Investigation, Formal analysis, Writing — review and editing. Vanesa Vinciauskaite: Resources, Investigation, Writing — review and editing. Marilyn Paul: Investigation, Formal Analysis. Rebecca Gilley: Investigation, Writing — review and editing. Simon Cook: Conceptualisation, Resources, Writing – review and editing. Glenn R Masson: Conceptualisation, Resources, Supervision, Investigation, Funding acquisition, Methodology, Writing — original draft, Project administration, Writing — review and editing.

## Notes

### Competing Interest Statement

The authors have declared no competing interest.

### Summary of Updates

The order of authors was incorrect. I, Glenn Masson, should be listed as the final author.

